# A spatially resolved mechanistic growth law for cancer drug development predicting tumour growing fractions

**DOI:** 10.1101/2021.05.03.442516

**Authors:** Adam Nasim, James Yates, Gianne Derks, Carina Dunlop

## Abstract

Mathematical models used in pre-clinical drug discovery tend to be empirical growth laws. Such models are well suited to fitting the data available, mostly longitudinal studies of tumour volume, however, they typically have little connection with the underlying physiological processes. This lack of a mechanistic underpinning restricts their flexibility and inhibits their direct translation across studies including from animal to human. Here we present a mathematical model describing tumour growth for the evaluation of single agent cytotoxic compounds that is based on mechanistic principles. The model can predict spatial distributions of cell subpopulations and account for spatial drug distribution effects within tumours. Importantly, we demonstrate the model can be reduced to a growth law similar in form to the ones currently implemented in pharmaceutical drug development for pre-clinical trials so that it can integrated into the current workflow. We validate this approach for both cell-derived xenograft (CDX) and patient-derived xenograft (PDX) data. This shows that our theoretical model fits as well as the best performing and most widely used models. However, in addition the model is also able to accurately predict the observed growing fraction of tumours. Our work opens up current pre-clinical modelling studies to also incorporating spatially resolved and multi-modal data without significant added complexity and creates the opportunity to improve translation and tumour response predictions.

## Introduction

Preclinical evaluation of drug efficacy plays a fundamental role in the development of oncological treatments, with the aim being to predict pharmacologically active drug concentrations and guide dose exploration in the clinic. Data for these studies comes from longitudinal measurements of tumour volume in animal models with specific targets investigated by the use of both cell-derived xenograft (CDX) and patient-derived xenografts (PDX) [1]. Central to these preclinical studies are *mathematical* models used to describe the tumour dynamics, fit the experimental data, and evaluate the anti-tumour effect. These models usually take the form of simple growth laws for tumour volume. There are many such models which can satisfactorily capture the dynamics [2, 3, 4, 5, 6], so that for any given data set it is in practice very difficult to distinguish between them [4, 5, 7, 8, 9]. This is compounded by the fact that typically these growth functions are purely empirical descriptions of the data, not founded in a mechanistic description of the physiological process.

The extension of growth law modelling away from purely phenomenological laws into mechanistic modelling would have significant advantages. It would enable better discrimination between models, could improve translational efficacy and be flexible enough to incorporate new multi-modal data sets as they come online. Indeed current pre-clinical growth laws lack the flexibility to account for the significant physiological processes that are being proven to be key, typically not even accounting for nutrient driven variations in growth fraction dynamics and hypoxia, [10, 11, 12, 13]. However, from a practical perspective, more complex models raise significant additional challenges. In general they have more parameters, which makes it challenging to parameterise them, with consequent difficulty with parameter identifiability, with model over-parameterisation a key issue.

Here, we present a mechanistic and spatial mathematical model for tumour growth based on a well-accepted mathematical framework coupling spatial diffusive processes to tumour growth [14]. Significantly, its spatial underpinning means that it predicts the size of the growing fraction of the tumour over time. Under the assumption that the tumour is spherical we show that this model can be expressed as a growth law of a similar type to that currently used in the pharmaceutical industry. It thus can be fitted and analysed using current industry-standard methods. We demonstrate the ease of use by comparison with a current commonly adopted growth law and validate both the models using standard methods on CDX and PDX data. Another advantage of the model is it can predict tumour growth fraction, which is validated by end point histology. The new framework offers further significant advantages such as being able to account for spatial gradients in drug distribution and inform better treatment design. Together these demonstrate that the model is fit-for-purpose to bridge the gap between emerging multi-modal data availability and the current pre-clinical trial workflow.

## Materials and Methods

### Mathematical models for tumour growth and treatment effect

Mathematical models as used in pre-clinical studies typically can be expressed as a rate equation for the tumour volume *V* (*t*) of the form

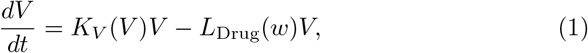

with the initial tumour volume *V* (*t* = 0) = *V*_0_, a fit parameter. The function *L*_Drug_(*w*) describes cell loss as a function of the drug concentration *w*(*t*) most often *L*_Drug_(*w*) = *K*_kill_*w*. The function *K*_*V*_ (*V*) represents the net growth rate, with a range of different forms used in pre-clinical studies [3, 5, 15] (Table 1) with the form often optimised for data fitting. For example, where growth is observed to be linear *K*_*V*_ (*V*) = *a/V*, whereas exponential growth is captured by *K*_*V*_ constant.

**Table 1:** Common growth models in pre-clinical modelling. The parameter *a* is the growth rate and *b* the rate of natural death. In the exponential-linear model *a*_0_ and *a*_1_ are the growth rates in the exponential and linear phases, respectively with *ψ* describing the transition between phases. The number of fit parameters for each model includes the initial tumour volume. (^*^: *ψ* = 20 is typically taken, ensuring a fast transition, so that it is effectively a three parameter system.)

| Model | $K_V$ | Parameters |
| --- | --- | --- |
| Linear | $\frac{a}{V}$ | 2 |
| Exponential | $a$ | 2 |
| Logistic | $a - bV$ | 3 |
| Gompertz | $a - b \ln(V)$ | 3 |
| Exponential-linear [2] | $\frac{a_0}{\left[1 + \left(\frac{a_0}{a_1} V\right)^\psi\right]^{\frac{1}{\psi}}}$ | 4* |
| Surface growth [16] | $aV^{-\frac{1}{3}}$ | 2 |
| Proliferative rim [17] | $a \left(1 - \left(1 - \frac{r_d}{\left(\frac{3V}{4\pi}\right)^{\frac{1}{3}}}\right)^3\right)$ | 3 |

An alternative approach to modelling tumour growth outside of pharmaceutical drug development has been to model the spatial diffusion of nutrients within tumours directly. It is assumed that nutrient availability determines the status of the cell, with cells experiencing low nutrient levels undergoing necrosis. This results in a map of the tumour with different sized cellular compartments for e.g. proliferative and non-proliferative necrotic compartments determined responsively from the environmental conditions [14, 18], see Fig. 1. This basic framework has been extensively built upon in the mathematical literature [19, 20]. The increasingly complex models are typically expressed in terms of partial differential equations which are either solved numerically themselves or coupled into finer scaled simulations.

**Figure 1:**
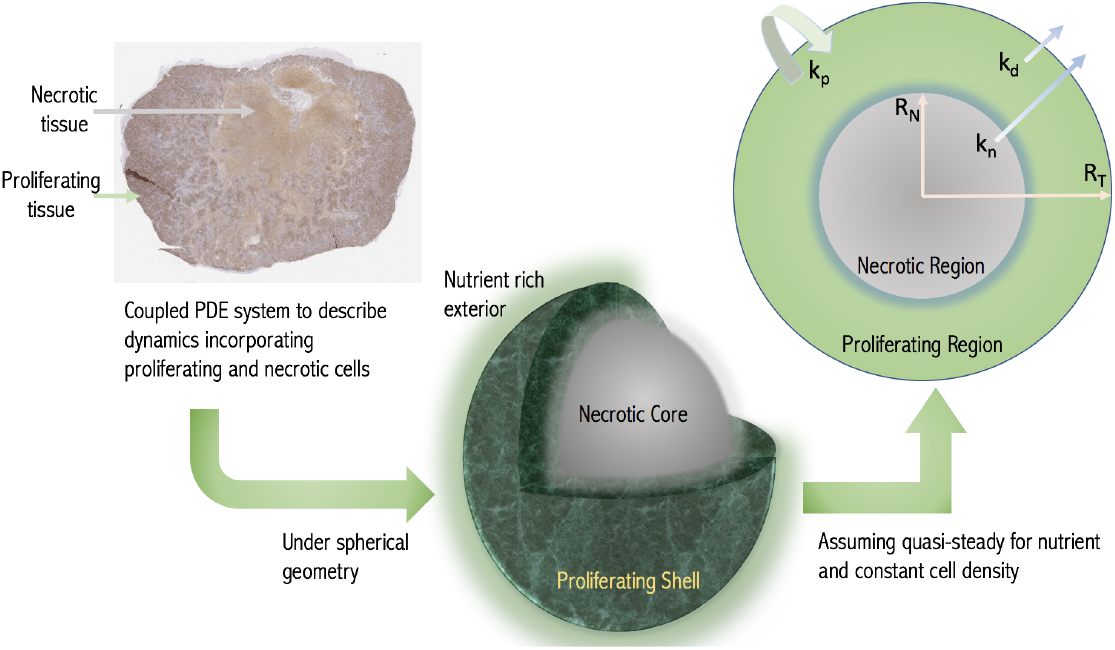
Schematic of the modelling concept. A growing tumour is modelled as a proliferating shell encapsulating a necrotic (non-proliferative core) with the boundaries between regions determined dynamically by considering nutrient diffusion. The assumed geometry and model variables and parameters are labelled in the cross-section with *R*_*N*_ and *R*_*T*_ being the necrotic and total tumour radii respectively. (Histological image - a day 24 CDX xenograft with Ki-67 stain.)

We adopt this diffusion-based framework but make simplifying assumptions appropriate to the pre-clinical context. We thus reduce the model to a growth law of the form Equation (1). The method is based on assuming a spherical tumour with the nutrient penetration within the tumour determined from the diffusion equation. Thus we obtain the growth fraction *GF* of the tumour, i.e. the volume fraction that is proliferating, with *GF* = 1 representing a fully proliferating tumour. A key parameter is the volume *V* ^*^ at which the tumour first experiences necrosis. The resulting model is similar to Equation (1), but with the growth fraction explicitly accounted for

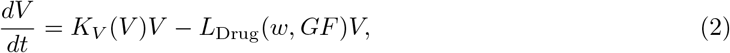

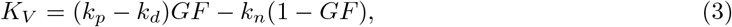

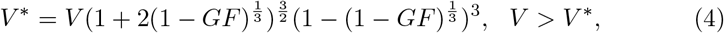

with *GF* = 1 when *V* < *V* ^*^ and *V* (0) = *V*_0_. In this model, *k*_*p*_ is the cell replication rate, *k*_*d*_ is the rate at which cells die in the proliferating region and *k*_*n*_ is the rate of breakdown and removal within the necrotic region. While the volume of the tumour *V* < *V* ^*^, the entire tumour proliferates and the model predicts exponential growth before switching to diffusion-limited growth as the volume increases with the growth fraction then determined from Equation (4). (See SI for further details).

The formulation Equations (2) to (4), which we term the diffusion-limited model is of the same mathematical form as currently implemented. However, it is now possible to directly predict the growth fraction of the tumour. We will demonstrate the functionality of the description Equations (2) to (4) both for control data sets and for drug trial data. We then take *L*_Drug_(*w, GF*) = *K*_kill_ *GF w*, which is superficially similar to the standard cell loss term, however loss now only occurs in the proliferating compartment. We see that the mechanistic approach allows us to incorporate that cell cycle specific pharmaceutical agents target proliferating cells. The PK description is in general compound specific. Here we consider CPT-11 (Irinotecan) which has been shown to be adequately described by an exponential model [13]. We thus set for the fitting *w* = *w*_0_ exp (−*αt*), with *w*_0_ the known administered dose and *α* the decay rate, leaving *K*_kill_ as the single parameter to be fitted describing drug effects. With the half life of Irinotecan being 12 hours, *α* = 1.39 day^−1^.

### Numerical implementation

Equations (2) to (4) form a growth law and two constraints which can be numerically solved as an algebraic differential equation. However, given current fitting protocols it is easier to work with a system of differential equations. This may be obtained from Equations (2) to (4) by differentiating the constraint Equation (4). The routines are implemented within MATLAB2019a (MathWorks, Natick, MA). Due to the significant inter-subject heterogeneity we use a non-linear mixed effects fitting approach (NLME) to fit both the population and individual subject parameters using the stochastic approximation expectation-maximization algorithm (SAEM). (See Supplementary Information).

### Data sets used for model validation

Control data are from AstraZeneca mouse-derived cell line xenografts (CDXs) for two cell-lines SW620 (29 mice), epithelial colorectal adenocarcinoma, and Calu6 (178 mice), epithelial lung adenocarcinoma. With regards treatment data we consider SW620 CDXs (95 mice) which were treated with weekly doses of 50mg/kg of CPT-11 (Irinotecan) either three times on days 1,8,15 (protocol 1, 68 mice), or four times during the experiment on days 1,8,15,22 (protocol 2, 18 mice) or 4,11,18,25 (protocol 3, 9 mice).

Additionally, we use the Novartis patient-derived xenograft (PDX) dataset, which is the largest publicly available database of PDX control data [21]. The Novartis data contains 226 mice with six different tumour types; breast carcinoma (BRCA) 39 mice, non-small cell lung carcinoma (NSCLC) 28 mice, gastric cancer (GC) 44 mice, colorectal cancer (CRC) 45 mice, cutaneous melanoma (CM) 33 mice, and pancreatic ductal carcinoma (PDAC) with 37 mice.

All animal studies in the United Kingdom were conducted in accordance with the UK Home Office legislation, the Animal Scientific Procedures Act 1986, and with AstraZeneca Global Bioethics Policy.

## Results

### Demonstration of the functionality of the model in fitting pre-clinical data

To demonstrate the fitting of the diffusion-limited model to longitudinal data we consider the AstraZeneca CDX data. In Fig. 2A–D we show the individual fits of the diffusion-limited model to the experimental data both combined and separated by protocol. The diffusion-limited model is observed to fit well across the range of curves. Similarly qualitatively good fits are observed for the other CDX and PDX control data sets (Fig. S.1). To further demonstrate the quality of fit, we perform a visual predictive check (VPC) [22]. This confirms that the diffusion-limited model captures the full range of dynamics (Fig. 2E and F). Similar results are also obtained from visual predictive checks for the other control PDX and CDX data sets (Fig. S.2).

**Figure 2:**
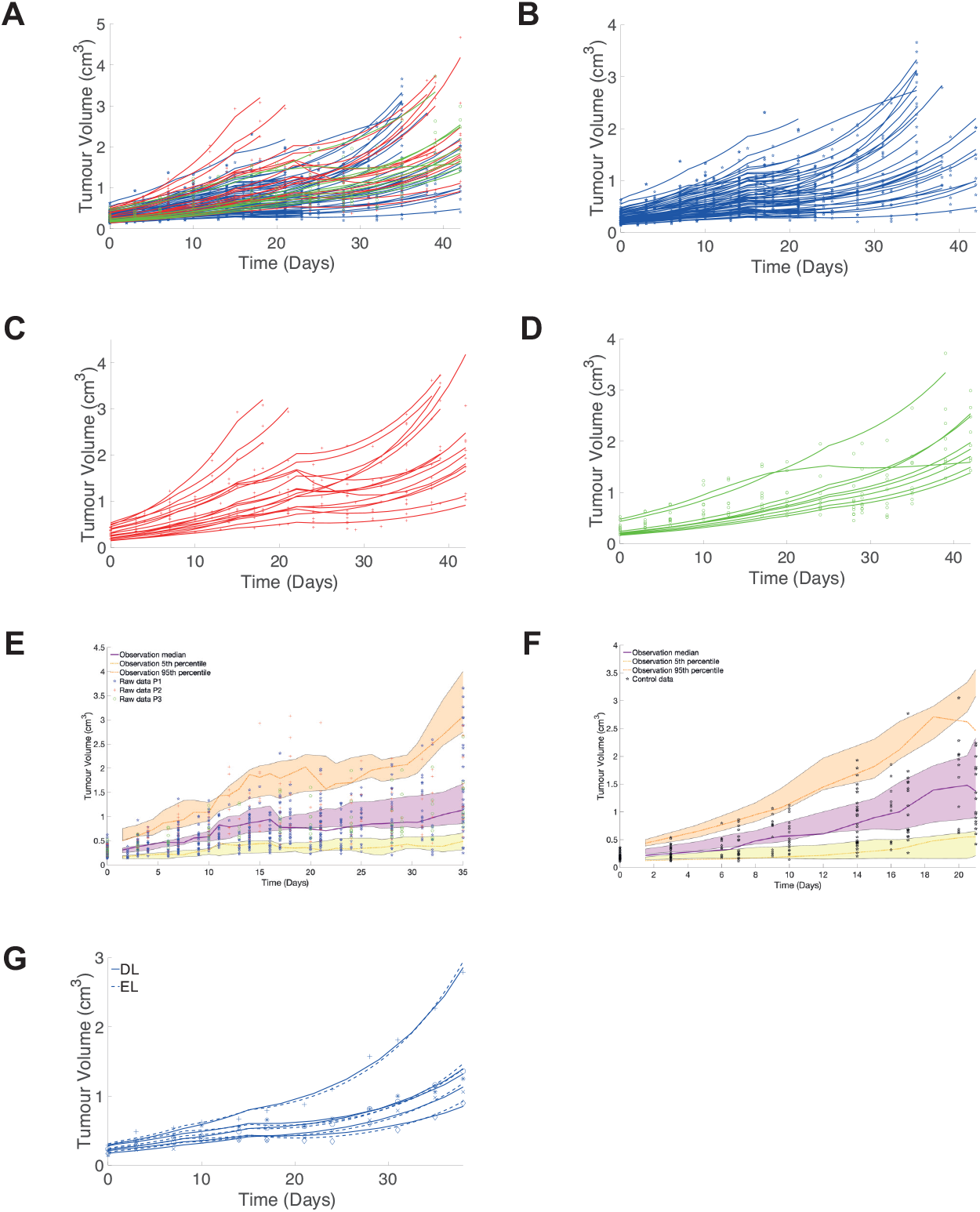
**A**, Fit of the diffusion-limited model to all CDX treatment data colour coded by treatment protocol (95 subjects, SW620 cell line, see Methods). **B**–**D**, Fits separated by treatment protocol for clarity. **E** and **F**, Visual predictive checks (VPC) of both the treated and control CDX data. The VPC is based on 1000 simulations, the shaded regions represent the 95% confidence intervals (CI) of the 25th, 50th and 95th percentiles of the simulated data. The experimental data median, 25th and 95th percentiles are marked (obtained using rolling average). **G**, Fit of both the diffusion-limited and exponential-linear models to five representative data sets from protocol 1.

Looking quantitatively at the NLME results (Table 2) we see a small residual mean square error. We further compare the diffusion-limited fit to results obtained from fitting one of the most widely used models, the exponential-linear model [2, 13, 23]. The parameter estimation for both the exponential-linear and diffusion-limited models is given in Table 2. Fig. 2G shows five representative fits. We clearly see the similarity in the model behaviour. This similarity in the quality of fit is confirmed by the closeness of the Akaike information criterion number (−217 and -221, for the diffusion-limited and exponential-linear respectively) and root mean square error (0.146 and 0.142, respectively) (Table 2). A similar result is achieved when considering the control data sets. These results are summarised in Tables S.1 to S.8. Given the typically limited time course of pre-clinical data it has been shown across numerous studies that no model may be optimal across all data sets [3, 5, 9]. However, the results indicate that the diffusion-limited model provides at least a similar standard of fit for typical data.

**Table 2:** NLME results of 10 runs for the full AstraZeneca CDX data set of 95 mice for the diffusion-limited (DL) and exponential-linear (EL) models.

| Model | Parameter | Definition | Value | SD | rmse | AIC |
| --- | --- | --- | --- | --- | --- | --- |
| DL | $k_p - k_d$ | Net growth rate(day<sup>-1</sup>) | 0.12 | 0.02 | 0.146 | -217 |
| | $k_n$ | Necrotic loss rate(day<sup>-1</sup>) | 0.05 | 0.03 | | |
| | $K_{\text{kill}}$ | Drug potency(kg mg<sup>-1</sup> day<sup>-1</sup>) | $5.7 \times 10^{-3}$ | $2.4 \times 10^{-4}$ | | |
| | $R^* (= (3V^*/4\pi)^{1/3})$ | Critical Radius(cm) | 0.41 | 0.07 | | |
| | $V_0$ | Initial Volume(cm<sup>3</sup>) | 0.25 | 0.11 | | |
| EL | $\lambda_0$ | Exponential growth rate(day<sup>-1</sup>) | 0.08 | 0.25 | 0.142 | -221 |
| | $\lambda_1$ | Linear growth rate(day<sup>-1</sup>) | 0.31 | 0.06 | | |
| | $\hat{k}(= k_1/30)$ | Transient death rate (day<sup>-1</sup>) | 24 | 6.2 | | |
| | $K_{\text{kill}}$ | Drug potency (kg mg<sup>-1</sup> day<sup>-1</sup>) | $1.2 \times 10^{-3}$ | $4 \times 10^{-4}$ | | |
| | $V_0$ | Initial Volume(cm<sup>3</sup>) | 0.29 | 0.13 | | |

### Diffusion-limited model predicts growth fraction dynamics with tumours

As discussed the rate equation predicts the growth fraction *GF* through Equation (4). Thus in addition to being able to fit the model to longitudinal measurements of tumour size we can now predict how the growth fraction changes, including in response to treatment. We show, for example, in Fig. 3A that those tumours showing greatest response (dosing strengths 75 and 100 mg/kg) are predicted to have the largest growth fraction, with consequent implications for treatment.

**Figure 3:**
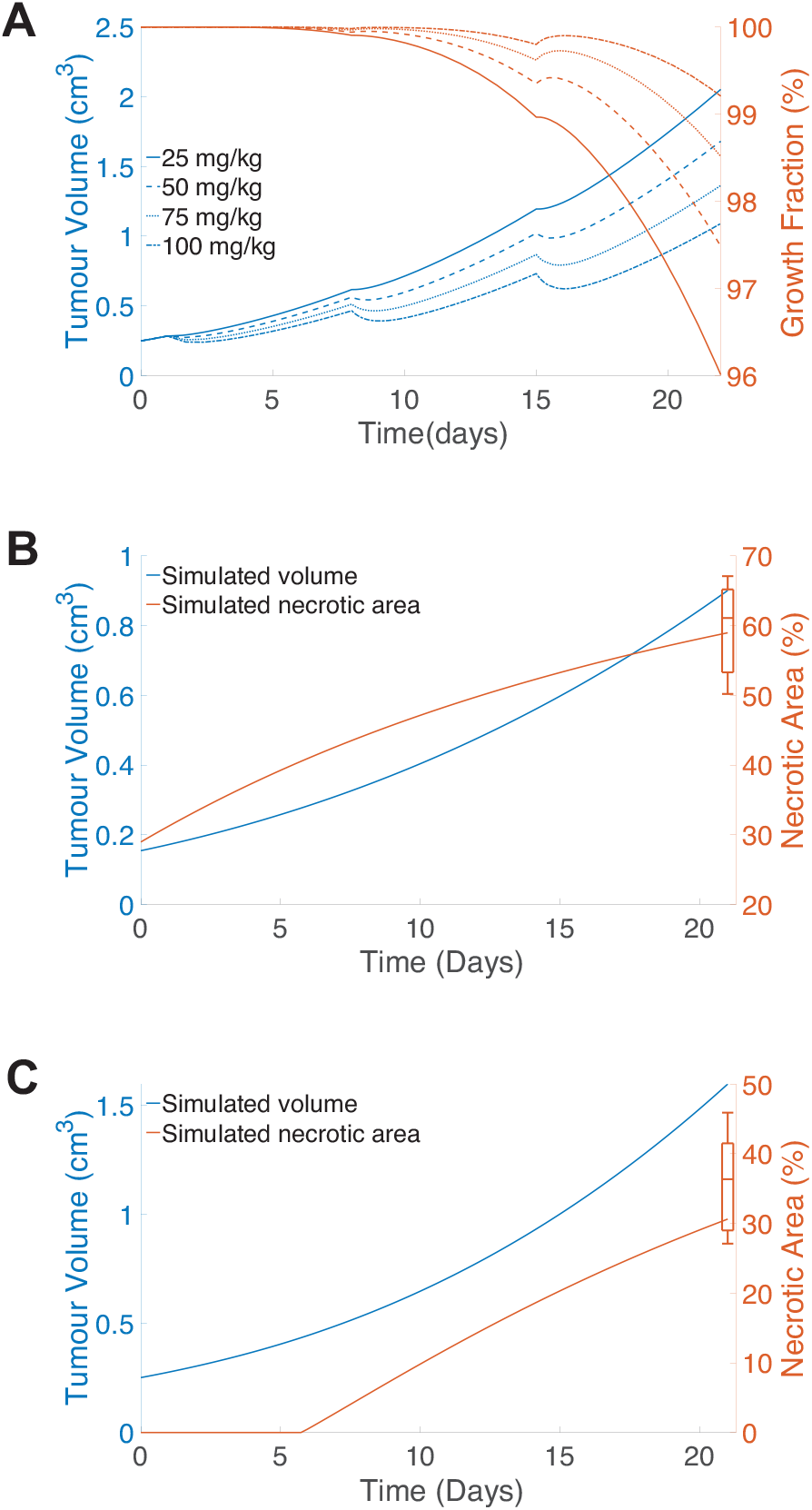
**A**, Simulated tumour response along with the predicted growth fraction for dosing strengths 25,50,75 and 100mg/kg dosed on days 1,8,15. **B**, and **C**, Simulated tumour dynamics for the CDX xenografts (**B**, SW620 and **C**, Calu6 cell lines, simulated curves using population parameters from Tables S.1 and S.2. The end-point box plots are derived from histological examination of necrotic area for 8 SW620 xenografts and 10 Calu6 xenografts.

The growth fraction of tumours is normally unavailable due to the difficulties of accessing this information *in vivo*. However, at the termination of xenograft experiments it is possible to obtain growth fraction data through histology, although this is not routinely done. We consider independent data sets for the necrotic area for untreated SW620 and Calu6 cell lines obtained from histology of bisected tumours (Ki-67 staining). The total volumetric growth fraction *GF* and necrotic area are related by 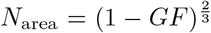. We simulate tumour dynamics for a tumour using the population parameters obtained from the mixed effects fitting of CDX data Tables S.1 and S.2. These simulations predict a necrotic area at the end of the experiment for both the SW620 cell line and the Calu6 cell lines that fits the experimental data (lying within the range of the data in Figs. 3B and C). In this case, the necrotic area data is an independent dataset. We could alternatively have used this data to further calibrate the model parameters. This highlights one of the benefits of a mechanistic model, that we can utilise other sources of information to better calibrate the model by way of multimodal fitting.

### Incorporation of spatial gradients in drug concentration

Not only spatial gradients of nutrients but also drug distribution can be incorporated in the model using the diffusion equation. This extended model, while more complex, is equally straightforward to numerically implement requiring a single additional integration. As a specific example, we consider the case where cell loss only occurs in proliferating cells. In this case, the cell loss is integrated over the proliferating compartment so that

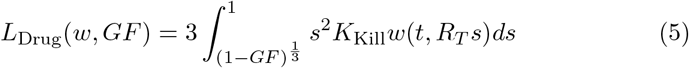

where *R*_*T*_ is the tumour radius, with for a spherical tumour *R*_*T*_ = (3*V/*4*π*)^1*/*3^. When the drug concentration *w*(*t*) is the same throughout the tumour we recover *L*_Drug_ = *K*_*Kill*_ *GF w* from Equation (5). For a drug diffusing into the tumour, however, the distribution is not constant and in this case the concentration in the proliferating region (where the drug acts) is (see Supplementary Information)

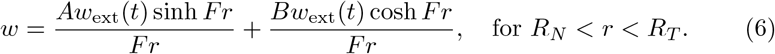

for *V* > *V* ^*^, where *w*_ext_(*t*) is the time varying applied drug concentration. The additional parameters *A*(*t*) and *B*(*t*) are known and defined in terms of *F, R*_*N*_, *R*_*T*_ and *D*_*w*_, see Supplementary Information. This reduces to 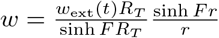 when *V* < *V* ^*^ and *GF* = 1. The parameter *F* effectively quantifies the depth of penetration of the drug with *F* = 0 corresponding to full drug distribution throughout the tumour. *F* large corresponds to a surface kill. In terms of the other parameters 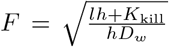, where *l* is the drug decay rate, *h* the drug efficiency and *D*_*w*_ the drug diffusivity.

The solution *w*(*r, t*) is then substituted into Equation (5) to determine *L*_Drug_. Although more complex than the previous model, it is easily implemented numerically. Heatmaps showing drug concentration within the tumour clearly demonstrate the effect of *F* (Fig. 4A). As *F* increases the amount of drug in the tumour drops significantly, indeed by the time *F* = 5 the concentration at the inner necrotic radius drops to 34% percent of its concentration at the outside edge (Fig. 4B). Exploring the effect of spatial drug distribution, we plot a representative solution of the spatial drug model (Fig. 4C). We see that as the drug penetration is reduced (*F* increased), the tumour displays more linear-like growth. With greater drug penetration (*F* small), a greater effect is observed with faster dynamic rebounds. Overall, it is clear tumour response will vary depending on the drug distribution.

**Figure 4:**
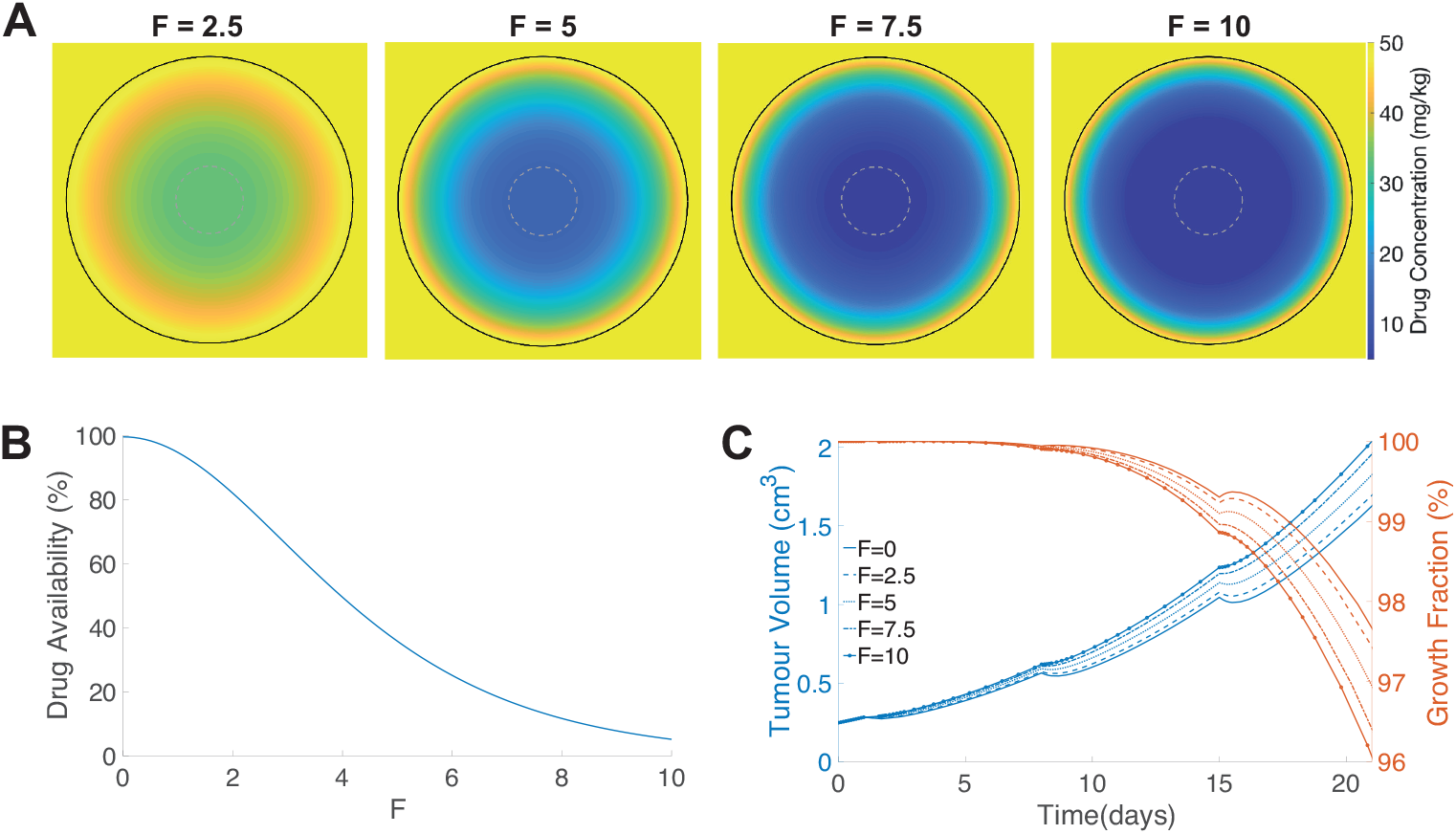
**A**, Heat map of the simulated internal drug distribution within the tumour immediately following dosing on day 15. The grey dashed circle shows the predicted necrotic region. The tumour edge is indicated by the black solid line. **B**, Percentage of drug concentration reaching inner necrotic radius (corresponding to grey circle in **A**) as a function of drug localization parameter *F*. **C**, Tumour volume and growth fraction dynamics for dosing strength fixed at 50mg/kg dosed on days 1,8,15. *F* = 0 describe full drug distribution, with *F* increasing corresponding to reducing drug penetration. At *F* = 10 the drug effect is largely restricted to the surface of the tumour.

## Discussion and Conclusions

We present a mathematical model for tumour growth and treatment that is based on a mechanistic description of nutrient limited growth. Despite being derived from spatial partial differential equations we show that these equations can be reduced to a growth law which is not significantly more complex than those currently used for pre-clinical drug trials.

The development of a mechanistic approach has several advantages over more phenomenological growth laws. Perhaps most significantly, in its current form the model allows for the dynamic prediction of the growing fraction of the tumour, accounting for cell loss from e.g. hypoxia. As most cytotoxic agents target only actively proliferating cells, tracking the growth fraction has potentially significant implications for treatment dynamics. This is only enhanced by the increasingly clear role hypoxia has in inducing downstream biological processes which directly promote tumour resistance to treatment [24, 25, 26, 27, 28].

By considering a range of different data sets including PDX and CDX data we have shown that the diffusion-limited model fits the data comparably to current industry-standard models. For independent datasets containing growth fraction data at termination we also predict a growth fraction commensurate with that observed. This without presenting significant increased complexity in the numerical solution. In addition, the diffusion-limited model parameters are all based on observable phenomena: net proliferation rate, maximum volume before necrosis begins, initial volume and cell loss through necrosis. The introduction of spatial modelling into a growth law framework has also enabled us to describe how the distribution of drug varies throughout the tumour. The indicative results obtained demonstrate the importance of accounting for this in pre-clinical studies. Indeed, the increasing importance of monoclonal antibody treatments (mAbs), with their reduced diffusivity, and antibody-drug conjugates (ADCs) [29] will only increase the importance of spatial modelling in pre-clinical trials.

## Acknowledgements

A. Nasim thanks the University of Surrey’s Doctoral College Grant and additionally AstraZeneca for funding. All authors thank AstraZeneca for generous access to data and the EPSRC funded QSP-UK network (EP/N005481/1) for facilitating initial discussions.

## Supplementary Information

### Tumour Growth Model Derivation

The paradigm diffusion-limited growth model of tumour growth [S1, S2, S3] is based on the solution of a diffusion model describing nutrient distribution within the tumour coupled to the equations of conservation of mass. We summarize the key points here.

The conservation of mass equation (with constant cell density) on the tumour domain is

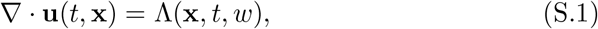

where **u** is the velocity field describing the motion of cells within the tumour and Λ is a growth function representing a mass source in proliferating regions and mass removal in necrotic regions. Cell proliferation increases the cellular mass which translates into tumour expansion and cell movement through the vector **u**. For the drug treatment models below Λ depends on the drug concentration *w* as well.

In conjunction with the conservation of mass equation we solve the diffusion equation for the concentration of nutrient in the tumour which is denoted by *c*. As nutrient diffusion is a fast process compared with tissue growth, the quasisteady approximation is valid leading to the diffusion equation in the form

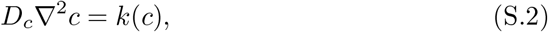

to be solved on the tumour domain, where *k*(*c*) is the cellular uptake rate. Assuming a critical concentration *c* = *c*_crit_ or equivalently a critical volume *V* ^*^ above which necrosis is initiated due to lack of nutrient, the cellular uptake is constant when *V* < *V* ^*^ but above this size a necrotic region forms so that cell uptake only occurs for growing cells.

After assuming radial symmetry, we introduce two dynamic radii:, the overall tumour radius *R*_*T*_ (*t*) and the radius at which the nutrient level drops low enough for necrosis to occur *R*_*N*_ (*t*) (only relevant if *R*_*T*_ corresponds to a volume larger than *V* ^*^), see Fig. 1. The radius *R*_*N*_ can also be expressed in terms of the growth fraction:

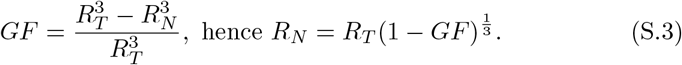

Under the assumption of spherical symmetry, the conservation of mass and diffusion equations reduce to

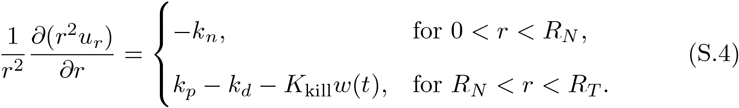

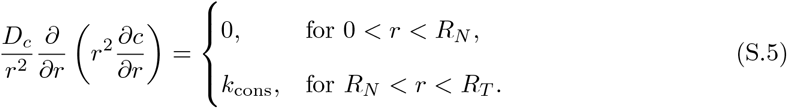

We can directly integratethe diffusion equation (S.5) on the domain *r* ≤ *R*_*T*_, with continuity conditions on *c* and *∂c/∂r* across the inner boundary, boundedness at *r* = 0. Two final conditions come from that at *r* = *R*_*T*_ the concentration is *c*_0_ which is the known concentration of nutrient outside the tumour and that at *R*_*N*_, *c* = *c*_crit_ which is the minimal concentration before necrosis occurs (which may alternatively be expressed in terms of *V* ^*^ the minimal volume). Importantly, evaluating the solution for concentration at *r* = *r*_*T*_ (where *c* = *c*_0_) determines the growth fraction as a function of the tumour volume through the constraint

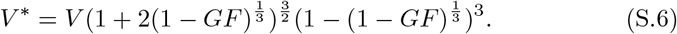

Finally, integrating the growth equation (S.4), in a similar manner gives a solution for the velocity **u**. Evaluating this solution on *r* = *r*_*T*_ where the velocity is 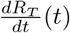 gives the growth law for the overall volume *V*, which after eliminating *R*_*N*_ using equation S.3 can be expressed in terms of the growth fraction as

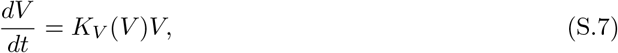

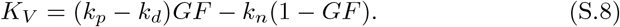

### Spatial drug distribution model

In a similar manner to the nutrient equation the spatial distribution of drug in the tumour is found by solving the diffusion equation for the drug concentration *w* on the tumour domain. We again assume a quasi-steady state due to drug diffusion being fast compared to tissue growth, a widely accepted assumption for cytotoxic chemotherapeutic agents [S4].

Assuming that drug uptake only occurs in the proliferative cells but with drug degradation continuing throughout the tumour, we get for the drug diffusion

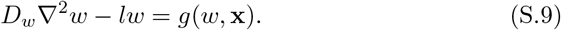

Here *D*_*w*_ is the diffusion constant of the particular compound in tumour tissue, *l* is the decay rate of the drug within the tumour. The function *g*(*w*, **x**) is the form for the drug action which we assume here to be linear 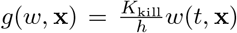, within the proliferation part of the tumour, where *K*_kill_ is the drug kill, and *h* > 0 is the efficiency of the drug, this is a measure of how much drug is used in the killing process: *h* → 0 indicates that large amounts of drug are required in the killing process whereas large *h* is the opposite. Finally *g*(*w*, **x**) is the functional form for the drug action which we assume here to be linear *g*(*w*, **x**) = *w*(*t*, **x**), within the proliferation part of the tumour. As for the nutrient equation, we obtain

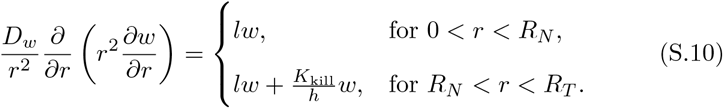

We require that the drug concentration as well as its derivative are continuous throughout the tumour domain as well as that the concentration is finite at the tumour centre, and denote the external drug concentration at the tumour by *w*(*t, r* = *R*_*T*_) = *w*_ext_(*t*), which is assumed known. This boundary value problem can be explicitly solved in terms of spherical harmonic functions [S5] which gives the distribution of drug concentration within the tumour when *V* > *V* ^*^ as

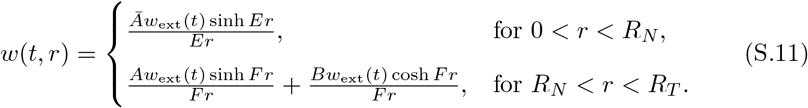

where 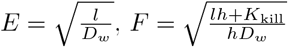. The parameters *Ā, A* and *B* are defined by

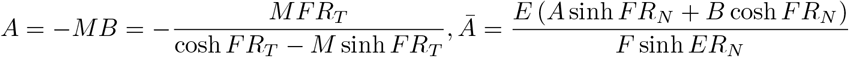

where 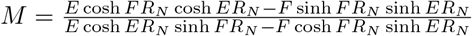. Note that as the drug kill effect needs only be considered in the proliferative compartment we only require the solution for *w*(*r, t*) only in *R*_*N*_ < *r* < *R*_*T*_. When *V* < *V* ^*^ and *GF* = 1 the solution reduces to 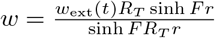.

Substituting for *w* from equation (S.11), proliferative compartment only, into the conservation of mass equation (S.4) and integrating as before gives the modified system

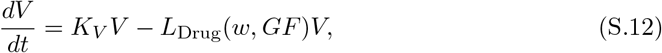

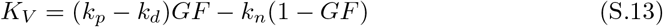

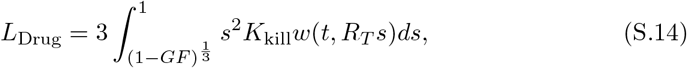

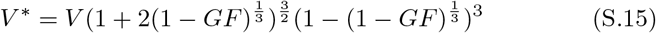

Note that if *F* = 0 (full drug distribution, hence *w*(*t, r*) = *w*_ext_(*t*)), the integral in (S.14) can be explicitly integrated and gives the relation for *K*_*V*_ as in (5).

### Numerical implementation

The diffusion limited model Equations (2) to (4) (main paper) may be implemented as an algebraic differential equation (ADE), however we choose to work with it as a system of ODEs. To this end we take the time derivative of the nutrient constraint Equation (4), this gives

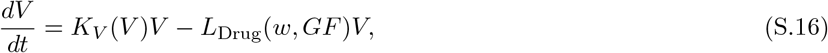

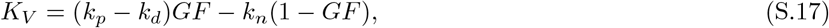

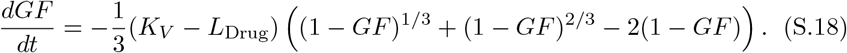

We solve (S.16)–(S.18) using MATLAB(2019a) using the ODE45 solver for the exponential phase and the ODE15S solver for the non-exponential phase using an event solver to manage the switch between phases. A stiff solver is required due to the high stiffness that may occur during integration from high dose strengths.

The mixed effects data fitting was implemented within MATLAB(2019a) using the NLMEFITSA optimisation routine. (With the necrotic radius *R*^*^ used as a fit parameter rather than *V* ^*^.) A linear approach to the approximation of the log likelihood was chosen. This was due to it being much faster to converge to a solution compared to other approximate techniques (Gaussian quadrature and Importance sampling were also tested, but they did not give significantly different results and took far longer to run). We used a probit transform for the parameter transform which is a natural way to bound the parameters (a logit transform could also have been used). The probit transform is defined as

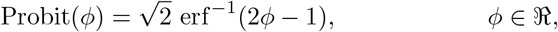

where erf is the error function. This probit transform is appropriate for all parameters except the rate of damage parameter *k*_1_ for the exponential-linear model. This parameter is required to be greater than one and so it was scaled as indicated in Table 2. Where standard deviations are calculated these are obtained from the individual parameters for each individual fit. These were transformed from the probit scale into the physical domain using the inverse probit transform and the standard deviation subsequently calculated.

---

**Table S.1:** NLME results of 10 runs for untreated SW620 CDX dataset 29 mice.

| Model | Parameter | Definition | Value | SD | rmse | AIC |
| --- | --- | --- | --- | --- | --- | --- |
| DL | $k_p - k_d$ | Net growth rate(day<sup>-1</sup>) | 0.131 | 0.021 | 0.086 | -150 |
| | $k_n$ | Necrotic loss rate (day<sup>-1</sup>) | 0.021 | 0.012 | | |
| | $R^*(= (3V^*/4\pi)^{1/3})$ | Critical Radius (cm) | 0.251 | 0.007 | | |
| | $V_0$ | Initial Volume (cm<sup>3</sup>) | 0.185 | 0.088 | | |
| EL | $\lambda_0$ | Exponential growth rate(day<sup>-1</sup>) | 0.103 | 0.062 | 0.090 | -139 |
| | $\lambda_1$ | Linear growth rate(day<sup>-1</sup>) | 0.157 | 0.327 | | |
| | $V_0$ | Initial Volume(cm<sup>3</sup>) | 0.182 | 0.321 | | |

**Table S.2:** NLME results of 10 runs for untreated Calu6 CDX dataset 178 mice.

| Model | Parameter | Definition | Value | SD | rmse | AIC |
| --- | --- | --- | --- | --- | --- | --- |
| DL | $k_p - k_d$ | Net growth rate(day<sup>-1</sup>) | 0.091 | 0.018 | 0.067 | -1193 |
| | $k_n$ | Necrotic loss rate (day<sup>-1</sup>) | 0.038 | 0.020 | | |
| | $R^*(= (3V^*/4\pi)^{1/3})$ | Critical Radius (cm) | 0.485 | 0.118 | | |
| | $V_0$ | Initial Volume (cm<sup>3</sup>) | 0.255 | 0.111 | | |
| EL | $\lambda_0$ | Exponential growth rate(day<sup>-1</sup>) | 0.086 | 0.025 | 0.072 | -1197 |
| | $\lambda_1$ | Linear growth rate(day<sup>-1</sup>) | 0.203 | 0.058 | | |
| | $V_0$ | Initial Volume(cm<sup>3</sup>) | 0.266 | 0.112 | | |

**Table S.3:** NLME results of 10 runs for untreated PDAC PDX dataset 37 mice.

| Model | Parameter | Definition | Value | SD | rmse | AIC |
| --- | --- | --- | --- | --- | --- | --- |
| DL | $k_p - k_d$ | Net growth rate(day<sup>-1</sup>) | 0.054 | 0.190 | 0.056 | -889 |
| | $k_n$ | Necrotic loss rate(day<sup>-1</sup>) | 0.055 | 0.041 | | |
| | $R^*(= (3V^*/4\pi)^{1/3})$ | Critical Radius (cm) | 0.458 | 0.086 | | |
| | $V_0$ | Initial Volume (cm<sup>3</sup>) | 0.259 | 0.035 | | |
| EL | $\lambda_0$ | Exponential growth rate(day<sup>-1</sup>) | 0.062 | 0.019 | 0.057 | -922 |
| | $\lambda_1$ | Linear growth rate(day<sup>-1</sup>) | 0.040 | 0.025 | | |
| | $V_0$ | Initial Volume(cm<sup>3</sup>) | 0.264 | 0.031 | | |

**Figure S.1:**
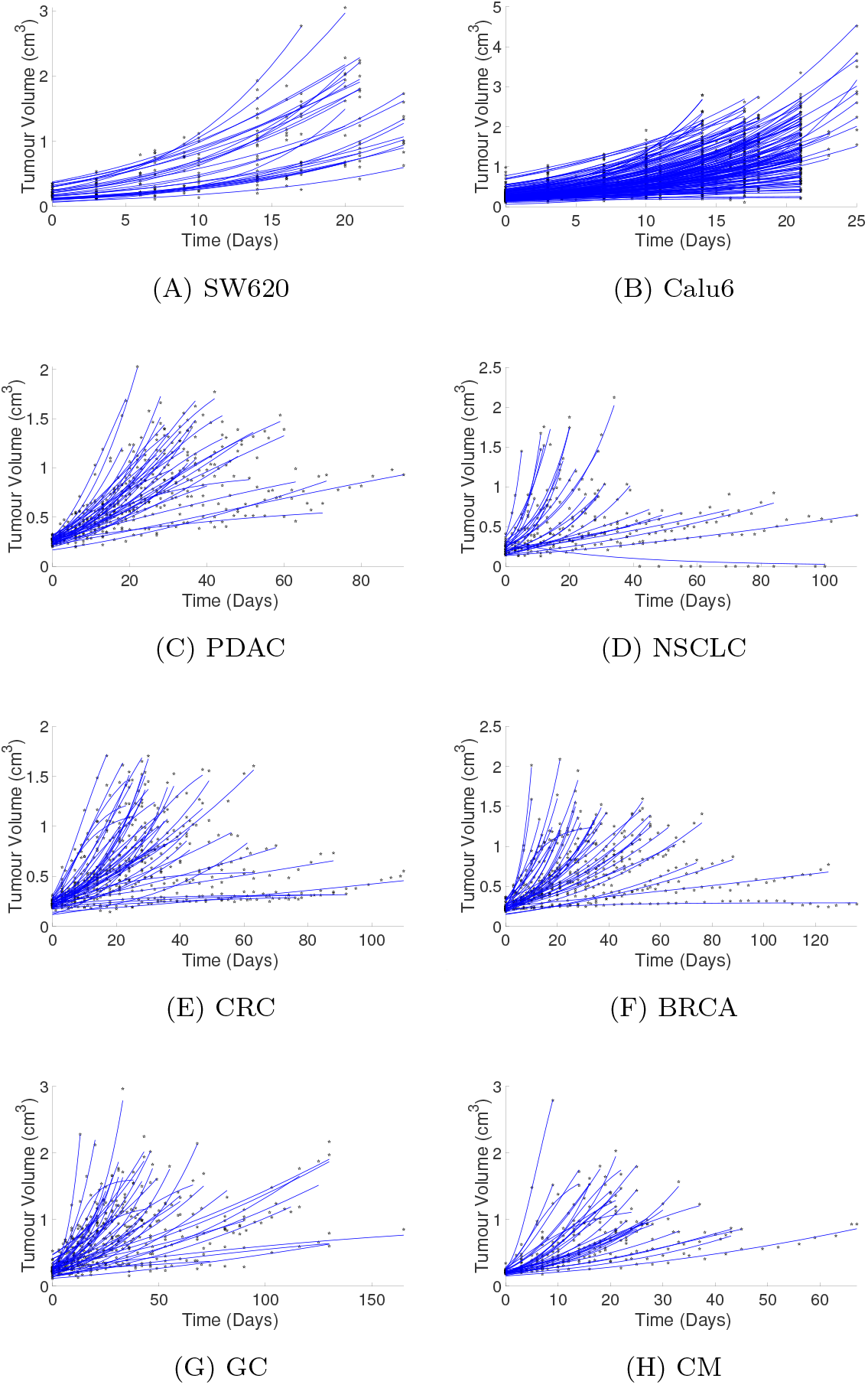
(a) to (h) show the fitted curves generated via the NLME optimisation for the diffusion limited model for all untreated data sets considered. (A) and (B) CDX control data sets from AstraZeneca for SW620 and Calu6 cell lines. (C)–(H) PDX control data sets from Novartis dataset [S6]. The stars indicate the volume data and the solid lines the simulated tumour growth curves.

**Figure S.2:**
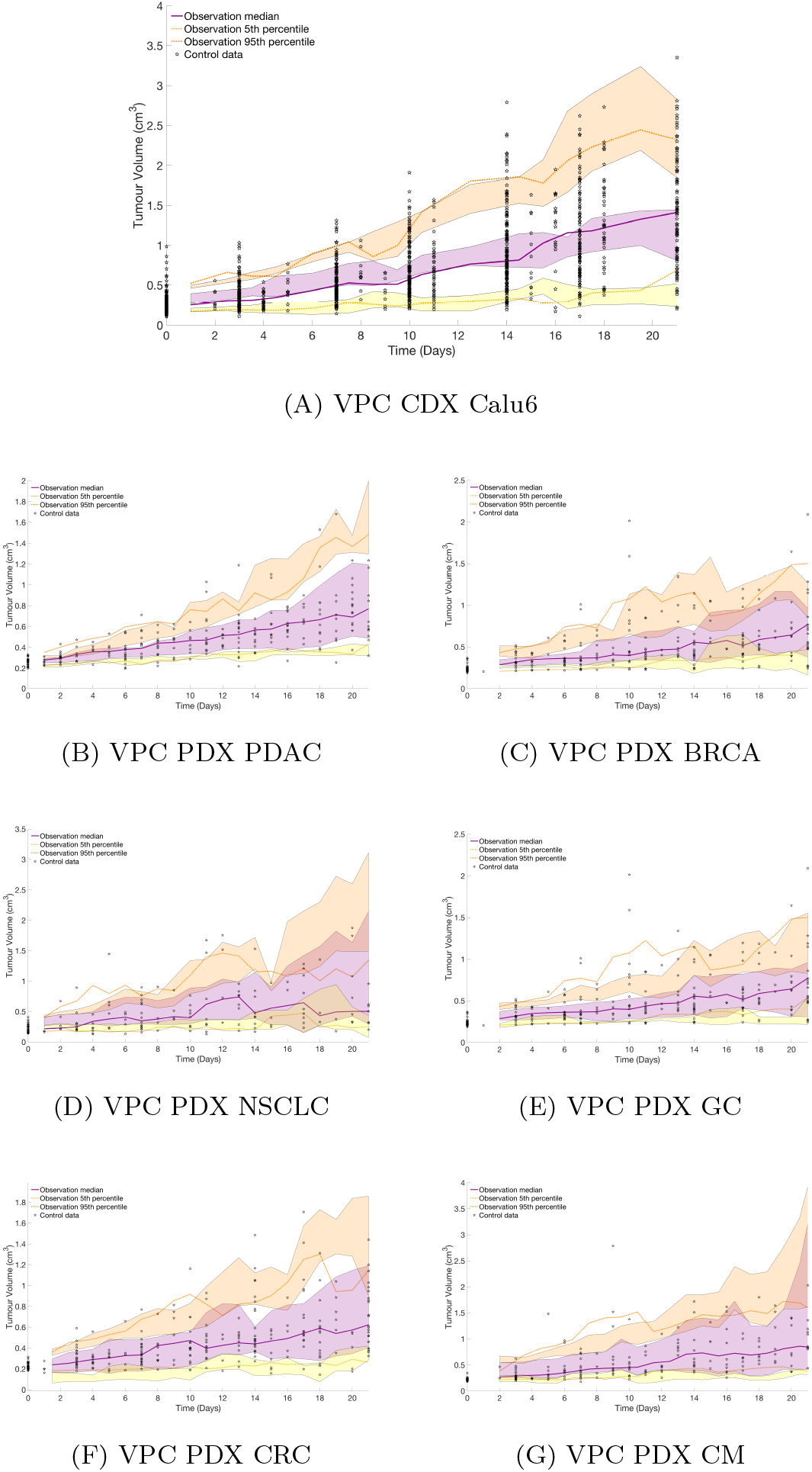
(A)–(G) shows the visual predictive check (VPC) for the control experiments considered in Fig. S.1. The VPC for the SW620 CDX data is found in the main paper. In all figures the orange, purple and yellow clouds indicate the 95th, 50th and 5th percentile confidence intervals respectively.

**Table S.4:** NLME results of 10 runs for untreated BRCA PDX dataset 39 mice.

| Model | Parameter | Definition | Value | SD | rmse | AIC |
| --- | --- | --- | --- | --- | --- | --- |
| DL | $k_p - k_d$ | Net growth rate(day<sup>-1</sup>) | 0.061 | 0.190 | 0.046 | -1096 |
| | $k_n$ | Necrotic loss rate(day<sup>-1</sup>) | 0.004 | $4 \times 10^{-4}$ | | |
| | $R^* (= (3V^*/4\pi)^{1/3})$ | Critical Radius (cm) | 0.6 | 0.086 | | |
| | $V_0$ | Initial Volume (cm<sup>3</sup>) | 0.247 | 0.035 | | |
| EL | $\lambda_0$ | Exponential growth rate(day<sup>-1</sup>) | 0.053 | 0.562 | 0.049 | -1075 |
| | $\lambda_1$ | Linear growth rate(day<sup>-1</sup>) | 0.183 | 3.153 | | |
| | $V_0$ | Initial Volume(cm<sup>3</sup>) | 0.246 | 0.054 | | |

**Table S.5:** NLME results of 10 runs for untreated NSCLC PDX dataset 28 mice.

| Model | Parameter | Definition | Value | SD | rmse | AIC |
| --- | --- | --- | --- | --- | --- | --- |
| DL | $k_p - k_d$ | Net growth rate(day<sup>-1</sup>) | 0.085 | 0.052 | 0.068 | -397 |
| | $k_n$ | Necrotic loss rate(day<sup>-1</sup>) | 0.010 | 0.004 | | |
| | $R^* (= (3V^*/4\pi)^{1/3})$ | Critical Radius(cm) | 0.385 | 0.19 | | |
| | $V_0$ | Initial Volume(cm<sup>3</sup>) | 0.218 | 0.068 | | |
| EL | $\lambda_0$ | Exponential growth rate(day<sup>-1</sup>) | 0.062 | 0.067 | 0.076 | -356 |
| | $\lambda_1$ | Linear growth rate(day<sup>-1</sup>) | 0.193 | 0.037 | | |
| | $V_0$ | Initial Volume(cm<sup>3</sup>) | 0.220 | 0.061 | | |

**Table S.6:**
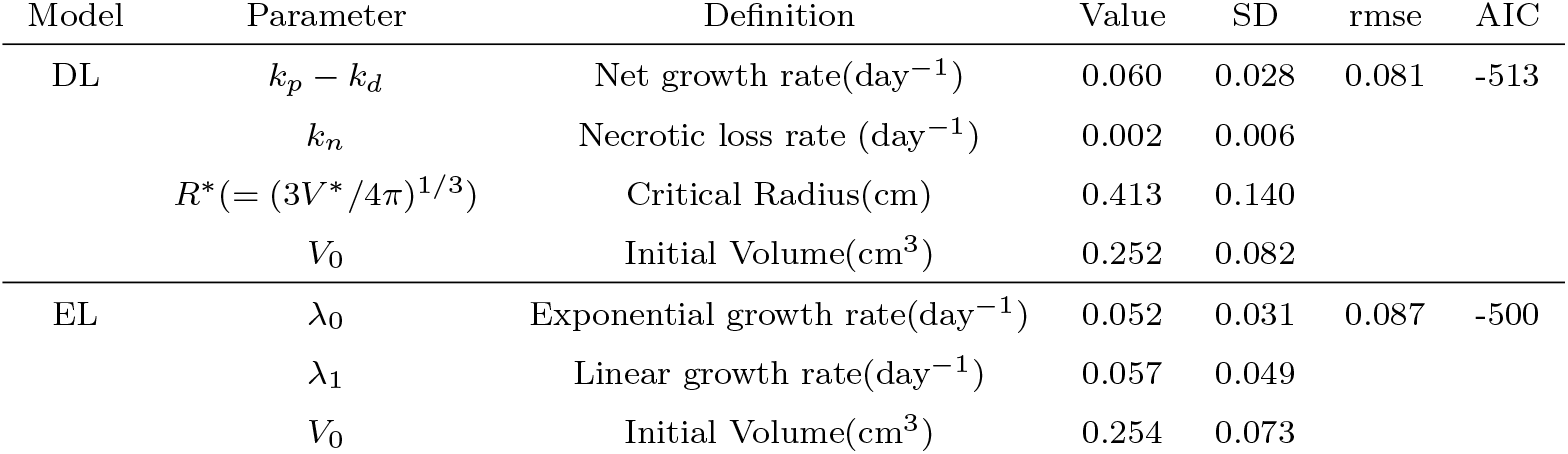
NLME results of 10 runs for untreated GC PDX dataset 44 mice.

| Model | Parameter | Definition | Value | SD | rmse | AIC |
| --- | --- | --- | --- | --- | --- | --- |
| DL | $k_p - k_d$ | Net growth rate(day<sup>-1</sup>) | 0.060 | 0.028 | 0.081 | -513 |
| | $k_n$ | Necrotic loss rate (day<sup>-1</sup>) | 0.002 | 0.006 | | |
| | $R^* (= (3V^*/4\pi)^{1/3})$ | Critical Radius(cm) | 0.413 | 0.140 | | |
| | $V_0$ | Initial Volume(cm<sup>3</sup>) | 0.252 | 0.082 | | |
| EL | $\lambda_0$ | Exponential growth rate(day<sup>-1</sup>) | 0.052 | 0.031 | 0.087 | -500 |
| | $\lambda_1$ | Linear growth rate(day<sup>-1</sup>) | 0.057 | 0.049 | | |
| | $V_0$ | Initial Volume(cm<sup>3</sup>) | 0.254 | 0.073 | | |

**Table S.7:**
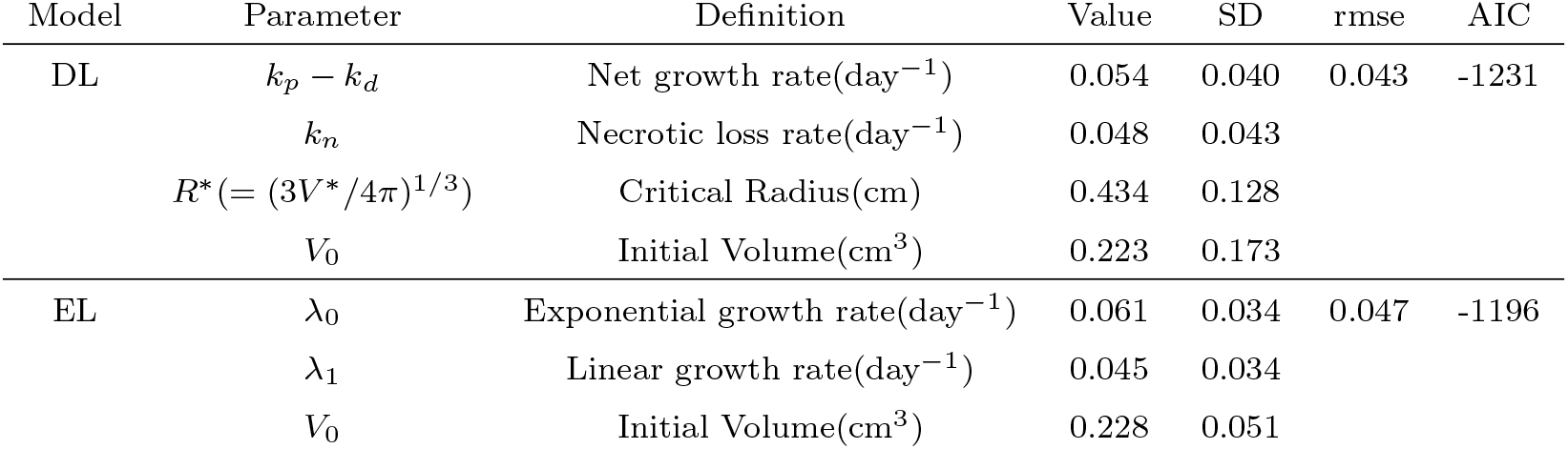
NLME results of 10 runs for untreated CRC PDX dataset 45 mice.

**Table S.8:**
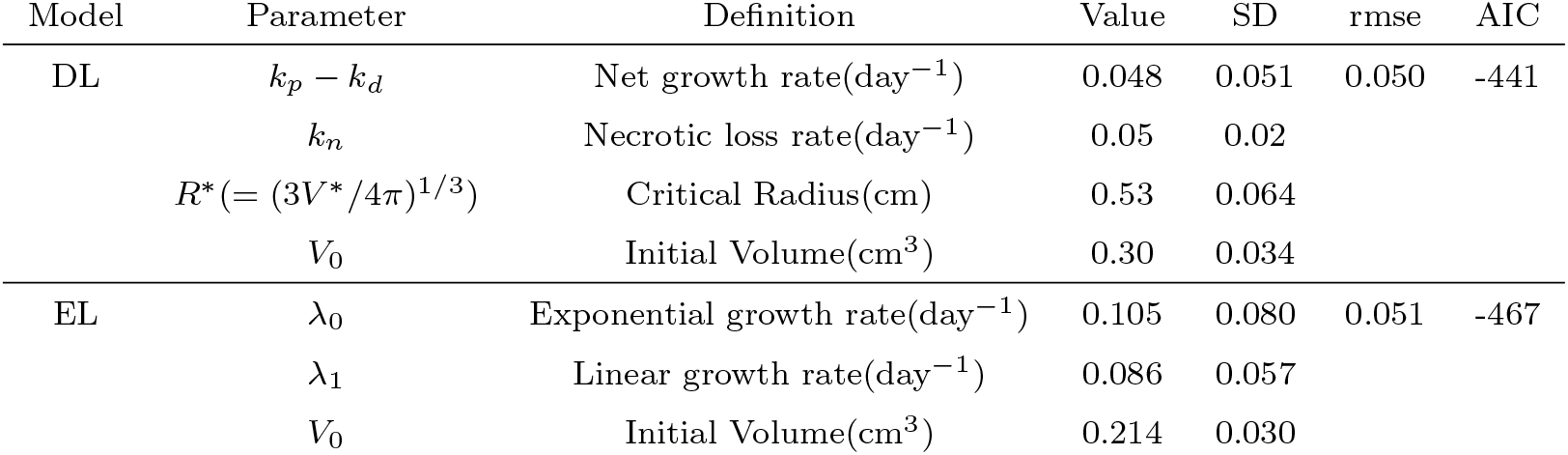
NLME results of 10 runs for untreated CM PDX dataset 33 mice.

| Model | Parameter | Definition | Value | SD | rmse | AIC |
| --- | --- | --- | --- | --- | --- | --- |
| DL | $k_p - k_d$ | Net growth rate(day<sup>-1</sup>) | 0.048 | 0.051 | 0.050 | -441 |
| | $k_n$ | Necrotic loss rate(day<sup>-1</sup>) | 0.05 | 0.02 | | |
| | $R^*(= (3V^*/4\pi)^{1/3})$ | Critical Radius(cm) | 0.53 | 0.064 | | |
| | $V_0$ | Initial Volume(cm<sup>3</sup>) | 0.30 | 0.034 | | |
| EL | $\lambda_0$ | Exponential growth rate(day<sup>-1</sup>) | 0.105 | 0.080 | 0.051 | -467 |
| | $\lambda_1$ | Linear growth rate(day<sup>-1</sup>) | 0.086 | 0.057 | | |
| | $V_0$ | Initial Volume(cm<sup>3</sup>) | 0.214 | 0.030 | | |

## References

[1] J.W.T Yates, H. Byrne, S.C. Chapman, T. Chen, J Delgado-SanMartin, G.D. Veroli, S. J. Dovedi, C. Dunlop, R. Jena, D. Jodrell, E. Martin, F. Mercier, A. Ramos-Montoya, H. Struemper, and P. Vicini. Opportunities for quantitative translational modeling in oncology. Clin. Pharmacol. Ther., 108(3):447–457, 2020.

[2] M. Simeoni, P. Magni, C. Cammia, G. De Nicolao, V. Croci, E. Pesenti, M. Germani, I. Poggesi, and M. Rocchetti. Predictive pharmacokineticpharmacodynamic modeling of tumor growth kinetics in xenograft models after administration of anticancer agents. Cancer Res., 64(3):1094–1101, 2004.

[3] B Ribba, NH Holford, P Magni, I Trocóniz, I Gueorguieva, P Girard, C Sarr, M Elishmereni, C Kloft, and LE. Friberg. A review of mixed-effects models of tumor growth and effects of anticancer drug treatment used in population analysis. CPT Pharmacometrics Syst., 3(5):113, 2014.

[4] H. Murphy, H. Jaafari, and H. Dobrovolny. Differences in predictions of ode models of tumor growth: A cautionary example. BMC cancer, 16:163, 2016.

[5] S. Benzekry, C. Lamont, A. Beheshti, A. Tracz, J. M. Ebos, L. Hlatky, and P. Hahnfeldt. Classical mathematical models for description and prediction of experimental tumor growth. PLoS Comput. Biol., 10(8):e1003800, 2014.

[6] E. Tjörve and K. M. Tjörve. A unified approach to the Richards-model family for use in growth analyses: why we need only two model forms. J. Theor. Biol., 267(3):417–425, 2010.

[7] M. Kühleitner, N. Brunner, W. G. Nowak, K. Renner-Martin, and K. Scheicher. Best fitting tumor growth models of the von Bertalanffy-PütterType. BMC Cancer, 19(1):683, 2019.

[8] D. Voulgarelis and J. Yates. NLME comparison of tumour growth models. Preprint., 2021.

[9] M. Marus̆ić, Z ̆. Bajzer, S. Vuk-Pavlović, and J.P. Freyer. Tumor growth in vivo and as multicellular spheroids compared by mathematical models. Bull. Math. Biol., 56(4):617–631, 1994. PMID: 15209404.

[10] X. Jing, F. Yang, C. Shao, K. Wei, M. Xie, H. Shen, and Y. Shu. Role of hypoxia in cancer therapy by regulating the tumor microenvironment. Mol Cancer, 18(1):157, 2019.

[11] M. A. Aleskandarany, A. R. Green, E. A. Rakha, R. A. Mohammed, S. E. Elsheikh, D. G. Powe, E. C. Paish, R. D. Macmillan, S. Chan, S. I. Ahmed, and I. O. Ellis. Growth fraction as a predictor of response to chemotherapy in node-negative breast cancer. Int J Cancer, 126(7):1761–1769, 2010.

[12] P. Gerlee. The model muddle: in search of tumor growth laws. Cancer Res., 73(8):2407–2411, 2013.

[13] S. Wilson, M. Tod, A. Ouerdani, A. Emde, Y. Yarden, A. Adda Berkane, S. Kassour, M. X. Wei, G. Freyer, B. You, E. Grenier, and B. Ribba. Modeling and predicting optimal treatment scheduling between the antiangiogenic drug sunitinib and irinotecan in preclinical settings. CPT Pharmacometrics Syst. Pharmacol., 4(12):720–727, 2015.

[14] H. M. Byrne and M. A. Chaplin. Growth of necrotic tumors in the presence and absence of inhibitors. Math. Biosci., 135(2):187–216, 1996.

[15] L. Carrara, S. M. Lavezzi, E. Borella, G. De Nicolao, P. Magni, and I. Poggesi. Current mathematical models for cancer drug discovery. Expert Opin. Drug Discov., 12(8):785–799, 2017.

[16] W. V. Mayneord. On a law of growth of Jensen’s rat sarcoma. Clin. Cancer Res., 16(4):841–846, 1932.

[17] N. D. Evans, R. J. Dimelow, and J. W. Yates. Modelling of tumour growth and cytotoxic effect of docetaxel in xenografts. Comput. Methods Programs Biomed., 114(3):3–13, 2014.

[18] N.F. Britton. Essential Mathematical Biology. Springer, 2005.

[19] J. S. Lowengrub, H. B. Frieboes, F. Jin, Y. L. Chuang, X. Li, P. Macklin, S. M. Wise, and V. Cristini. Nonlinear modelling of cancer: bridging the gap between cells and tumours. Nonlinearity, 23(1):R1–R9, 2010.

[20] H. M. Byrne. Dissecting cancer through mathematics: from the cell to the animal model. Nat. Rev. Cancer, 10(3):221–230, 2010.

[21] H. Gao, J. M. Korn, S. Ferretti, J. E. Monahan, Y. Wang, M. Singh, C. Zhang, C. Schnell, G. Yang, Y. Zhang, O. A. Balbin, S. Barbe, H. Cai, F. Casey, S. Chatterjee, D. Y. Chiang, S. Chuai, S. M. Cogan, S. D. Collins, E. Dammassa, N. Ebel, M. Embry, J. Green, A. Kauffmann, C. Kowal, R. J. Leary, J. Lehar, Y. Liang, A. Loo, E. Lorenzana, E. Robert McDonald, M. E. McLaughlin, J. Merkin, R. Meyer, T. L. Naylor, M. Patawaran, A. Reddy, C. Röelli, D. A. Ruddy, F. Salangsang, F. Santacroce, A. P. Singh, Y. Tang, W. Tinetto, S. Tobler, R. Velazquez, K. Venkatesan, F. Von Arx, H. Q. Wang, Z. Wang, M. Wiesmann, D. Wyss, F. Xu, H. Bitter, P. Atadja, E. Lees, F. Hofmann, E. Li, N. Keen, R. Cozens, M. R. Jensen, N. K. Pryer, J. A. Williams, and W. R. Sellers. High-throughput screening using patientderived tumor xenografts to predict clinical trial drug response. Nat. Med., 21(11):1318–1325, 2015.

[22] M. Bergstrand, A. C. Hooker, J. E. Wallin, and M. O. Karlsson. Prediction-corrected visual predictive checks for diagnosing nonlinear mixed-effects models. AAPS J, 13(2):143–151, 2011.

[23] N. Terranova and P. Magni. TGI-Simulator: a visual tool to support the preclinical phase of the drug discovery process by assessing in silico the effect of an anticancer drug. Computat Methods Programs Biomed, 105(2):162–174, 2012.

[24] S. Riffle and R. S. Hegde. Modeling tumor cell adaptations to hypoxia in multicellular tumor spheroids. J. Exp. Clin. Cancer Res., 36(1):102, 2017.

[25] J. M. Brown and W. R. Wilson. Exploiting tumour hypoxia in cancer treatment. Nat. Rev. Cancer, 4(6):437–447, 2004.

[26] S. Strese, M. Fryknäs, R. Larsson, and J. Gullbo. Effects of hypoxia on human cancer cell line chemosensitivity. BMC Cancer, 13:331, 2013.

[27] L. Zheng, C. J. Kelly, and S. P. Colgan. Physiologic hypoxia and oxygen homeostasis in the healthy intestine. A review in the theme: Cellular responses to hypoxia. Am. J. Physiol. Cell Physiol., 309(6):C350–360, 2015.

[28] S. Daster, N. Amatruda, D. Calabrese, R. Ivanek, E. Turrini, R. A. Droeser, P. Zajac, C. Fimognari, G. C. Spagnoli, G. Iezzi, V. Mele, and M. G. Muraro. Induction of hypoxia and necrosis in multicellular tumor spheroids is associated with resistance to chemotherapy treatment. Oncotarget, 8(1):1725–1736, 2017.

[29] V. S. Goldmacher and Y. V. Kovtun. Antibody-drug conjugates: using monoclonal antibodies for delivery of cytotoxic payloads to cancer cells. Ther. Deliv., 2(3):397–416, 2011.

## References

[S1] J. S. Lowengrub, H. B. Frieboes, F. Jin, Y. L. Chuang, X. Li, P. Macklin, S. M. Wise, and V. Cristini. Nonlinear modelling of cancer: bridging the gap between cells and tumours. Nonlinearity, 23(1):R1–R9, 2010.

[S2] H. M. Byrne and M. A. Chaplin. Growth of necrotic tumors in the presence and absence of inhibitors. Math. Biosci., 135(2):187–216, 1996.

[S3] N.F. Britton. Essential Mathematical Biology. Springer, 2005.

[S4] V. A. Levin, C. S. Patlak, and H. D. Landahl. Heuristic modeling of drug delivery to malignant brain tumors. J. Appl. Biopharm., 8(3):257–296, 1980.

[S5] A. Abramowitz and I.A. Stegun. Pocketbook of Mathematical Functions. Harri Deutsch Verlag, 1984.

[S6] H. Gao, J. M. Korn, S. Ferretti, J. E. Monahan, Y. Wang, M. Singh, C. Zhang, C. Schnell, G. Yang, Y. Zhang, O. A. Balbin, S. Barbe, H. Cai, F. Casey, S. Chatterjee, D. Y. Chiang, S. Chuai, S. M. Cogan, S. D. Collins, E. Dammassa, N. Ebel, M. Embry, J. Green, A. Kauffmann, C. Kowal, R. J. Leary, J. Lehar, Y. Liang, A. Loo, E. Lorenzana, E. Robert McDonald, M. E. McLaughlin, J. Merkin, R. Meyer, T. L. Naylor, M. Patawaran, A. Reddy, C. Röelli, D. A. Ruddy, F. Salangsang, F. Santacroce, A. P. Singh, Y. Tang, W. Tinetto, S. Tobler, R. Velazquez, K. Venkatesan, F. Von Arx, H. Q. Wang, Z. Wang, M. Wiesmann, D. Wyss, F. Xu, H. Bitter, P. Atadja, E. Lees, F. Hofmann, E. Li, N. Keen, R. Cozens, M. R. Jensen, N. K. Pryer, J. A. Williams, and W. R. Sellers. High-throughput screening using patient-derived tumor xenografts to predict clinical trial drug response. Nat. Med., 21(11):1318–1325, 2015.

